# Sex and Circuit Specific Dopamine Transporter Regulation Underlies Unique Behavioral Trajectories of Functional *SLC6A3* Coding Variation

**DOI:** 10.1101/2021.11.02.466932

**Authors:** Adele Stewart, Felix P. Mayer, Raajaram Gowrishankar, Gwynne L. Davis, Lorena B. Areal, Paul J. Gresch, Rania M. Katamish, Rodeania Peart, Samantha E. Stilley, Keeley Spiess, Maximilian J. Rabil, Faakhira A. Diljohn, Angelica E. Wiggins, Roxanne A. Vaughan, Maureen K. Hahn, Randy D. Blakely

**Affiliations:** Department of Biomedical Science, Florida Atlantic University, Jupiter, FL; Wilkes Honors College, Florida Atlantic University, Jupiter, FL; Brain Institute, Florida Atlantic University, Jupiter, FL; Department of Biomedical Sciences, University of North Dakota School of Medicine and Health Sciences, Grand Forks, North Dakota

## Abstract

Virtually all neuropsychiatric disorders display sex differences in prevalence, age of onset, and/or clinical symptomology. In sex-biased disorders, one sex is often suggested to harbor protective mechanisms, rendering them resilient to genetic and/or environmental risk factors. Here, we demonstrate sex-biased molecular, pharmacological and behavioral effects induced by the dopamine (DA) transporter (DAT) coding variant Ala559Val, previously identified in subjects diagnosed with the male-biased disorders attention-deficit/hyperactivity disorder and autism spectrum spectrum disorder. In DAT Val559 mice, we identified sex differences in response to psychostimulants, social behavior, and cognitive traits. We reveal a sex by circuit dissociation in D2-type autoreceptor (D2AR) regulation of DAT wherein D2AR-dependent DAT phosphorylation and trafficking, detectable in the male dorsal striatum, does not occur in females but rather is a property of the ventral striatum, predicting sex-specific changes in behavior. Consequently, we found that a subset of altered behaviors can be normalized using the D2R antagonist sulpiride in DAT Val559 mice. Our studies provide a cogent example of how sex shapes the behavioral trajectory of DA signaling perturbations and identify the sex-dependent, locality-selective capacity for D2AR regulation of DAT as an unrecognized determinant of this trajectory. Rather than identifying one sex as resilient, we find that sex can drive alterative behavioral patterns from shared signaling perturbations that may result in females being underreported. Our work underscores the utility of model systems to study the functional intrusions of rare genetic variation to gain insights into pathways underlying normal and perturbed trait domains associated with common neuropsychiatric conditions.

## Introduction

Signaling actions of the neurotransmitter dopamine (DA) mediate multiple complex behaviors including motor control, motivation, attention, response to novelty, reward, and cognition. Two DA synthesizing cell clusters in the midbrain innervate distinct areas of the forebrain. DA neurons of the substantia nigra pars compacta (SNc) project to the dorsal striatum (dStr), a critical component of the basal ganglia motor loop. DA neurons of the ventral tegmental area (VTA) project to the ventral striatum (vStr), comprised principally of the nucleus accumbens (NAc), which modulates circuits linked to reward prediction and response. The VTA also innervates the pre-frontal cortex (PFC), constituting the mesocorticolimbic projection, with activity linked to working memory and goal-driven behavior. Though DA neurons constitute a comparatively small population, dysfunctions in DA signaling have been implicated in a number of neurologic and neuropsychiatric disorders including Parkinson’s disease, attention-deficit/hyperactivity disorder (ADHD), schizophrenia, bipolar disorder (BPD), autism spectrum disorder (ASD) and addiction, each of which displays sex bias in prevalence, age of onset, and/or clinical symptomology^1-4^. Nonetheless, preclinical research targeting the functional and behavioral output of disease-linked disruptions in DA systems have predominantly been confined to the study of male animals^5^.

The presynaptic dopamine transporter (DAT, *SLC6A3*) mediates high-affinity clearance of DA and is the principal mechanism responsible for rapid extracellular DA inactivation in both dStr and vStr^6-8^. A growing body of evidence has linked female steroid hormones, particularly estradiol, to the control of DAT expression and function in rodents. Voltametry studies have confirmed that estradiol modulates striatal DA clearance^9^ and that clearance rates vary across the estrus cycle^10^. Similarly, DAT activity is greater in females^11^, and pharmacological disruption of either DAT or D2-type autoreceptor (D2AR) results in exaggerated changes in DA dynamics in the dStr of females^12^. More recently, Calipari et al. found that phosphorylation of DAT at Thr53, a site implicated in DAT activity and psychostimulant binding^13^, changes with estrus cycle stage, providing a molecular mechanism that can link the actions of estradiol to capacity for efficient DA signaling^10^. Non-hormonal mechanisms may also underlie sex differences in the DA system as DA release and reuptake have been found to differ across striatal subregions in slice preparations irrespective of estrus stage^14^.

While acknowledging the biological differences between the rodent and human brain, studies in animal models represent an important opportunity to explore biological mechanisms that contribute to sex bias of relevance for neuropsychiatric disease in an environment that lacks interference from sociocultural factors that influence outcomes in humans^15, 16^. To identify penetrant genetic changes in DA signaling that drive neuropsychiatric disease risk, and thereby generate improved, construct-valid animal models, we screened subjects with ADHD, a disorder treated with DAT-targeted psychostimulants including amphetamine (AMPH, Adderall®) and methylphenidate (MPH, Ritalin®), for evidence of functional DAT coding variation. Among the variants identified^17-19^, a Ala559Val substitution was found in two brothers^19^. The Ala559Val variant was was previously detected in a girl with BPD^20^ and subsequently identified in two unrelated boys with ASD^21^. *In vitro* studies revealed that the Val559 substitution promotes a form of nonvesicular, transporter-mediated DA release we designate as anomalous DA efflux (ADE)^22^. Male mice harboring the DAT Val559 mutation were found to display constitutive D2-type autoreceptor (D2AR) activation that suppresses vesicular DA release, reductions in both AMPH- and MPH-induced elevations in extracellular DA and hyperactivity^23^, waiting impulsivity and heightened motivation^24^. Recent work in the DAT Val559 model demonstrated D2AR-induced DAT phosphorylation and trafficking is confined to DAergic projections to the dStr, biasing DAT Val559-mediated perturbations in DA clearance to the nigrostriatal circuit^25^. Whether these changes were also observed in female animals was not examined, owing largely to the male bias of the disorders where DAT Val559 had been identified, though DAT Val559 was also found in a girl with a BPD diagnosis^20^, a disorder where females display differences in age of onset and clinical features that warrant distinct treatment considerations^26, 27^. In a search for endogenous factors that might confer resilience to DAT Va559 actions, we uncovered compelling evidence for sex-specific traits arising from expression of DAT Val559, including a sexually dimorphic capacity of D2ARs to regulate DAT that distinguishes nigrostriatal vs mesolimbic DA circuits.

## Materials and Methods

### Animals

All experiments utilizing animals were performed under a protocol approved by the Institutional Animal Care and Use Committee (IACUC) at either Vanderbilt University or Florida Atlantic University. Behavioral and surgical experiments were performed on wildtype (WT) and homozygous DAT Val559 littermate mice bred from heterozygous breeders maintained on a hybrid background (75% 129S6/SvEvTac and 25% C57BL/6J)^23^ Due to the association of the DAT Val559 mutation with neurodevelopmental disorders^19, 21^, behavioral testing was performed on juvenile/young adult animals 6-9 weeks of age. All animals were housed on a 12:12 light/dark cycle with water and food available *ad libitum*, and, unless otherwise noted, experiments were performed during the active phase. Due to technical limitations, animals utilized for microdialysis and amperometry experiments were between 8-12 weeks of age and experiments were performed during the inactive phase. For slice preparations, 4-6 week-old mice homozygous for DAT Val559 or WT mice were bred from homozygous dams and sires derived from heterozygous breeders, no more than two generations removed.

### Behavioral Assays

Behavioral testing was completed in either the Laboratory for Neurobehavior Core Facility operated by the Vanderbilt Brain Institute or in the Neurobehavior Core Facility at the Florida Atlantic University Stiles-Nicholson Brain Institute. Experiments were performed during the active phase of the light/dark cycle under red light. Animals were habituated to the testing rooms for a minimum of 20 min prior to the start of each experiment. All behavioral assays were performed by an experimenter blinded to animal genotype. For all experiments data are combined from at least two independent animal cohorts. Locomotor testing and conditioned place preference were conducted according to previously published protocols^23, 28^. Drugs were dissolved in sterile saline [D-Amphetamine hemisulfate salt, Sigma (3 mg/kg); methylphenidate HCl, Sigma (10 mg/kg); (±)-sulpiride, Sigma (50 mg/kg); SKF83822 hydrobromide, Tocris (0.5 mg/kg)] and administered via intraperitoneal injection (i.p.). Additional details regarding specific behavioral tests can be found in the Supplemental Materials and Methods.

### AMPH-induced striatal ERK 1/2 phosphorylation

Animals were separated into singly housed cages (day 1) and received saline injections on days 2 and 3 to habituate to injection stress. On day 4 animals received a single amphetamine injection (3 mg/kg) and rapid decapitation occurred 15 min post-drug injection. Brains were rapidly dissected on a pre-chilled metal stage and whole striata removed for homogenization in pre-heated (95°C) lysis buffer (1% SDS, 0.8 mM EDTA, 0.5 M Tris Base with protease and phosphatase inhibitors) with a hand-held homogenizer. Samples were placed on a 95°C heat block for 10 min, vortexed and spun at 10,500 rpm for 10 min. Supernatant was removed and processed for immunoblotting. Membranes were blocked for 1 hour at RT in a phosphoprotein blocker (Millipore) and probed for total (Cell Signaling, Danvers, MA, RRID:AB_10695739; 1:1000) and phospho-ERK 1/2 (Cell Signaling, RRID:AB_331646; 1:1000).). Li-cor Odyssey IRDye secondary antibodies were used (IRDye800 goat anti-mouse, 1:10,000; IRDye680 goat anti-rabbit, 1:15,000) to allow for multiplexing imaging of the protein bands. Phospho-ERK1/2 bands were normalized to their Total ERK1/2 bands to account for differences in loading.

### Acute brain slice studies

Brain slice preparation and experiments were performed under constant oxygenation (95%O_2_:5%CO_2_). Animals were killed by rapid decapitation and excised brains were moved to ice-cold sucrose-artificial CSF (S-ACSF: 250 mM sucrose, 2.5 mM KCl, 1.2 mM NaH_2_PO_4_, 26 mM NaHCO_3_, 11 mM D-glucose, 1.2 mM MgCl_2_, 2.4 mM CaCl_2_, pH 7.4, 300–310 mOsm). Brain slices containing dStr or vStr were prepared exactly as we previously described^25^. Before drug treatments, slices were washed with ACSF (substituting 92 mM NaCl for sucrose (pH 7.4, 300 – 310 mOsm)) at 37°C and hemisections obtained from the same slice were treated with either vehicle or 1 μM (-)-Quinpirole HCl (Sigma, St. Louis, MO) for 5 min (biotinylation) or 10 min (immunoprecipitation).

### Brain slice biotinylation studies

Acute slices were treated with sulfoNHS-SS-biotin (1 mg/ml, ThermoFisher, Waltham, MA) on ice for 30 min. Reactions were quenched by treatment with 0.1 M glycine 2 X for 10 min each, followed by 3 rapid and two 5-minute washes with ice-cold ACSF. dStr and vStr were dissected and tissue solubilized in radioimmunoprecipitation assay (RIPA) buffer, supplemented with protease inhibitor cocktail (SIGMA; P8340). Lysates were cleared of cellular debris by centrifugation (20,000 x g for 15 min at 4°C) and cleared supernatants were exposed to streptavidin agarose beads (ThermoFisher) at a ratio of 20 g protein per 50 μl bead slurry and mixed overnight at 4°C. Following washes with RIPA buffer, protein was eluted/denatured at room temp for 30 min and subjected to SDS-PAGE, transferred onto PVDF membranes, and immunoblotted with primary rat anti-DAT (MAB369, Millipore; RRID:AB_2190413; 1:1000) overnight at 4°C followed by the addition of a goat anti-rat HRP-conjugated secondary antibody (Jackson ImmunoResearch, Westgrove, PA; 1:10,000) for 1 hr. Chemiluminescence (Bio-Rad Clarity ECL, Hercules, CA) was visualized using an ImageQuant LAS 4000 imager (GE Healthcare Life Sciences, Marlborough, MA). Densitometric quantification of immunoblots was performed, and surface DAT levels were normalized to total DAT.

### DAT Thr53 phosphorylation studies

Lysates for immunoprecipitation of total and pThr53-DAT were prepared as above. Rabbit anti-Thr53 DAT antibody (RRID:AB_2492078^13^) was crosslinked to protein A magnetic beads (Dynabeads, ThermoFisher) as previously described^25^. Brain slices were solubilized in lysis buffer (in mM: NaCl 150, KCl 2.5 m, Tris 50, and 1% Triton X-100; supplemented with protease inhibitor cocaktail (P8340) and phosphatase inhibitor cocktail 3(Milipore Sigma, P0044)), cleared of cellular debris by centrifugation (20,000 x g for 15 min at 4°C) and cleared supernatants were added to cross-linked DAT Thr53 antibody-conjugated beads at a ratio of 250 μg protein to 25 μl of bead slurry, rocked at 4°C for 4 h, washed with lysis buffer, treated with 2 X Laemmli sample buffer (Bio-Rad) to elute/denature protein before SDS-PAGE and immunoblotting for DAT as above. Densitometric quantification of immunoblots was performed, and DAT pThr53 levels were normalized to total DAT.

### High-speed chronoamperometry

High-speed chronoamperometry (HSC) for DA was performed as described previously^29^. Peak DA signal amplitudes were generated in triplicate for each animal. The clearance rate (T_C_ in nM/s) was estimated as the slope of the linear portion of current decay from 20 to 60% signal amplitude. Michaelis Menten analyses were performed to obtain K_m_ values.

### Statistical Analyses

Data are presented as mean ± S.E.M and were analyzed using Prism 9.0 (GraphPad Inc, La Jolla, CA). Statistical significance was set at *P* < 0.05 for all analyses. A Student’s unpaired, two-sided t-test was used to compare between two independent data sets. For data containing two independent variables, two-way analysis of variance (ANOVA) was performed followed by Sidak’s *post hoc* test. Repeated measures ANOVA (rmANOVA) was utilized with Sidak’s *post hoc* tests for time course data sets. Uptake inhibition best-fit curves were generated using the log(inhibitor) vs. response standard curve function in Prism, allowing for variable slope. Tube test contingency tables were analyzed with Fisher’s exact test.

## Results

### Sex and DAT Val559 genotype influence post-synaptic mechanisms recruited following AMPH exposure

DAT Val559 females exhibited no alterations in baseline locomotion (Fig. 1a, S1a, S1b). However, whereas male DAT Val559 mice exhibit blunted AMPH-induced hyperactivity (3mg/kg i.p., Fig. 1b) relative to WT littermates, females, who demonstrated a reduced motor activation to AMPH compared to males, showed no genotype effect on AMPH motor responses (Fig. 1b, S1c, S1d). Male DAT Val559 mice also display blunted locomotor responses to the DAT blocker methylphenidate (MPH)^23^. Like with AMPH, DAT Val559 females exhibited no change in MPH (10 mg/kg, i.p.)-induced increases in distance traveled (Fig. S2a), stereotypy (Fig. S2b) or rearing (Fig. S2c). The ability of AMPH to inhibit DA uptake in striatal synaptosomes did not significantly differ between genotypes in either males or females (Fig. S3a). Similarly, the ability of AMPH to elevate extracellular DA was unaltered in DAT Val559 males (Fig. S3b), suggesting that alterations in presynaptic actions of AMPH on DA release cannot explain effects of the drug in mutant males. Interestingly, dStr microdialysis studies in DAT Val559 females indicated that AMPH-dependent increases in extracellular DA were blunted (Fig. S3b). No sex or genotype effects were seen in AMPH-stimulated 5-HT elevations (Fig. S3c). These findings pointed to the possibility that postsynaptic changes in DA receptor signaling might contribute to the sex-dependent locomotor actions of AMPH in DAT Val559 mice.

**Fig. 1.**
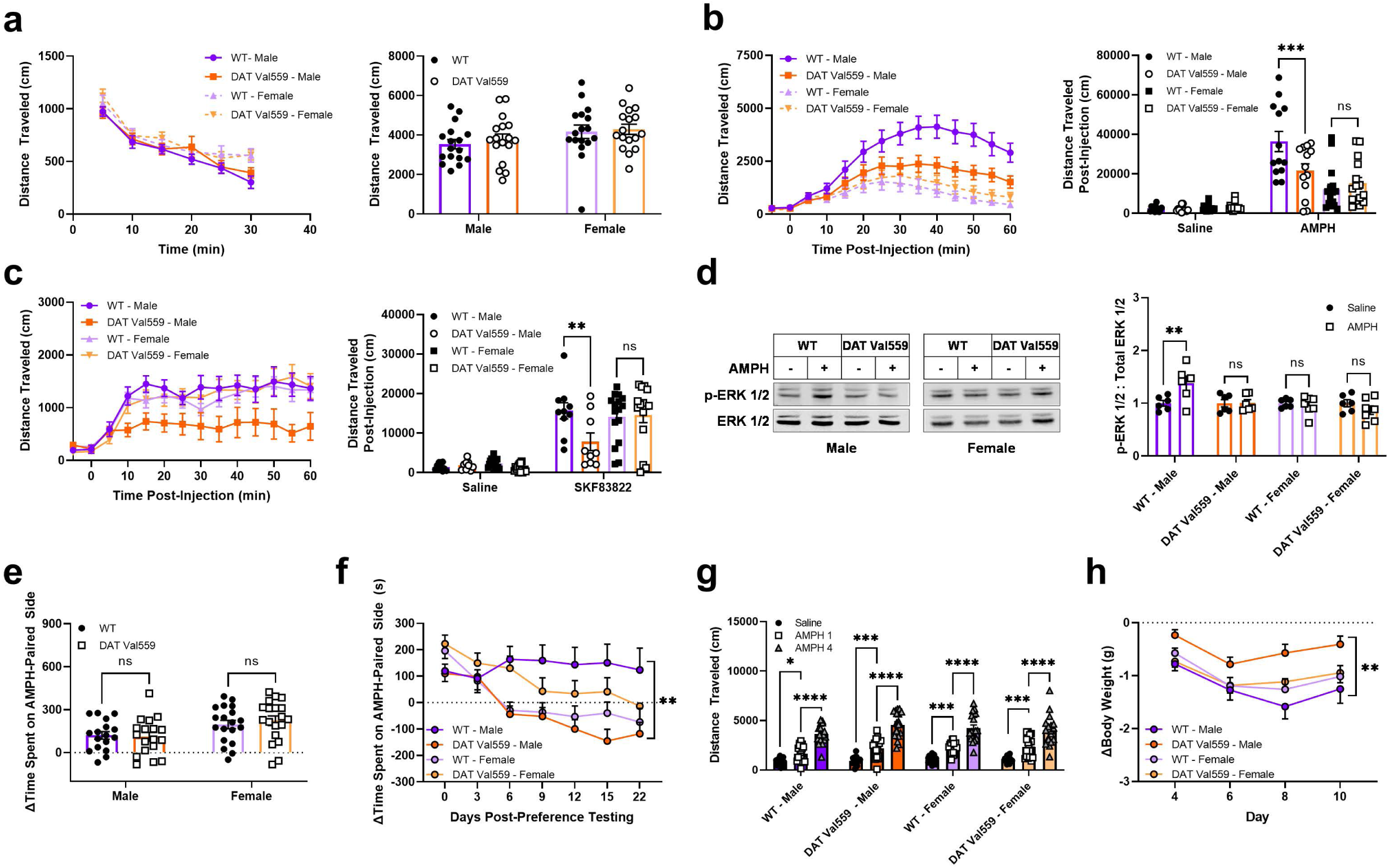
Male, but not female, DAT Val559 mice display blunted AMPH-dependent locomotor hyperactivity, accelerated CPP extinction, and blunted AMPH hypophagia. **a** WT (n=17) and DAT Val559 (n=17) male and females were allowed to freely explore an open field chamber for 30 mins. Distance traveled was monitored over time and summarized for the complete recording period. **b** Acute AMPH (3 mg/kg, i.p.)-driven horizontal distance traveled in WT male (n=13), WT female (n=16), DAT Val559 male (n=13) and DAT Val559 female (n=15) mice. Summary data for the 60 min recording period post-drug injection is provided. **c** WT male (n=10), WT female (n=16), DAT Val559 male (n=9) and DAT Val559 female (n=15) were administered a single injection of the D1R agonist SKF83822 (0.5 mg/kg, i.p.). Summary data for the 60 min recording period post-drug injection is provided. **d** Phosphorylation of ERK 1/2 was determined in whole striatal lysates from WT and DAT Val559 male (n=6) and female (n=6-7) mice following AMPH (3 mg/kg, i.p.) challenge (15 min). Representative immunoblots are shown. Phosphorylated ERK 1/2 was normalized to total protein and data are expressed relative to WT saline treated animals. AMPH (3 mg/kg, i.p.) CPP was performed in WT male (n=18), WT female (n=19), DAT Val559 male (n=18) and DAT Val559 female (n=19) mice. **e** CPP scores following 4 paired saline or AMPH injections. **f** Time course of AMPH CPP extinction. **g** AMPH locomotor sensitization. Data from saline day 1, AMPH day 1 and AMPH day 4 are depicted to highlight locomotor sensitization. **h** Body weight loss during chronic AMPH exposure. Data were analyzed by two-way ANOVA or two-way repeated measures ANOVA (time courses) with Sidak’s multiple comparisons test. ^*^*P*<0.05, ^**^*P*<0.01, ^***^*P*<0.001, ^****^*P*<0.0001. ns = not significantly different. Data are presented as mean ± SEM.

Post-synaptic actions of DA in the Str are mediated by D1 and D2 receptors (D1R and D2R), and as postsynaptically acting D2R agonists are unavailable, we turned to an examination of D1Rs. At high doses (2 mg/kg, i.p.), the ability of the D1R-selective agonist SKF83822 to stimulate locomotor behaviors is equivalent in WT and DAT Val559 males^23^. However, at a lower dose (0.5 mg/kg, i.p.), SKF83822-induced blunted horizontal locomotion (Fig. 1c) and stereotypy (Fig. S4a) in DAT Val559 males relative to WT males, as seen for AMPH, but not in females. Females did, however, exhibit decreased rearing (Fig. S4b) and center occupancy (Fig. S4c) at the same dose of SKF83822, suggesting D1R contribution to female phenotypes, though occurring in different circuits. Expression of D1Rs and D2Rs is unaltered in the striatum of male DAT Val559 mice^23^, leading us to postulate that loss of D1R-driven locomotor activation might result from receptor desensitization or uncoupling of key receptor effectors. To address this idea, we examined AMPH triggered phosphorylation of the critical signal transducer ERK 1/2. Systemic AMPH elevated ERK1/2 phosphorylation in WT, but not DAT Val559 males (Fig. 1d). Female WT mice, in contrast, did not display an elevation of ERK1/2 phosphorylation, despite displaying extracellular DA elevations comparable to that shown in males, nor was ERK1/2 phosphorylation elevated in DAT Val559 females. These data identify loss of DA-dependent ERK 1/2 activation as a possible mechanism contributing to blunting of D1R- and AMPH-dependent locomotor activation in DAT Val559 males.

### Impact of sex and DAT Val559 genotype on psychostimulant place preference and sensitization

To investigate whether sex dependence modulates the impact of DAT Val559 on the rewarding properties of AMPH we employed a conditioned place preference (CPP) paradigm to detect both the acquisition and extinction of drug-context memory. While female sex was associated with overall higher AMPH CPP scores (*P*=0.0026), neither genotype nor interaction effects were detected in the ability of animals to acquire AMPH CPP (Fig. 1e). However, while WT males failed to extinguish place preference out to 22 days (Fig. S5a), DAT Val559 males demonstrated extinction after 6 days. Female WT animals also reached a zero-preference index at 6 days, but AMPH CPP extinction in female DAT Val559 occurred at 9 days, a shift in the direction opposite to the genotype effect observed in males (Fig. 1f). We also assessed locomotor sensitization to repeated AMPH^30^, and found that, although DAT Val559 males, display blunted responses to the acute locomotor stimulating actions of AMPH (Fig. 1b)^23^, AMPH locomotor sensitization showed no genotype or sex differences (Fig. 1g, S5b). AMPH is also a potent appetite suppressant^31^. Indeed, the anorexigenic actions of AMPH were blunted in DAT Val559 males compared to WT animals, whereas females displayed no genotype effect (Fig. 1h).

### DAT Val559 mice exhibit sex-specific, non-motor behavioral phenotypes

The apparent “resilience” of DAT Val559 females to changes in psychostimulant-dependent phenotypes led us to hypothesize that female DAT Val559 mice might lack behavioral phenotypes evident in males. Prior work in DAT Val559 males detected no open space avoidance in the elevated zero maze^23^. Similarly, baseline center occupancy in the open field dose not differ between WT and DAT Val559 mice of either sex (Fig. S6a). However, we did note both sex and genotype effects on center exploration in response to psychostimulants. While MPH fails to promote center occupancy in mice of either sex or genotype (Fig. S6c), AMPH increased center exploration with a sex-dependence, occurring exclusively in WT and DAT Val559 males (Fig. S6b). These observations prompted our further examination of harm avoidance in DAT Val559 mice using the light/dark transition test, believed to exploit the conflict between a rodent’s propensity to explore new environments versus innate threat aversion. As shown in Fig. 2a, DAT Val559 males displayed an increase in the latency to enter the light compartment compared to WT littermates, though surprisingly they exhibited a significantly longer light side occupancy resulting in a longer latency to return to the dark side. Moreover, DAT Val559 males displayed fewer side transitions without a change in distance traveled (Fig. 2a). Female DAT Val559 mice, conversely, display a more typical “risk-averse” phenotype in this task with longer latencies to enter the light side and a decrease in light side occupancy (Fig. 2b).

**Fig. 2.**
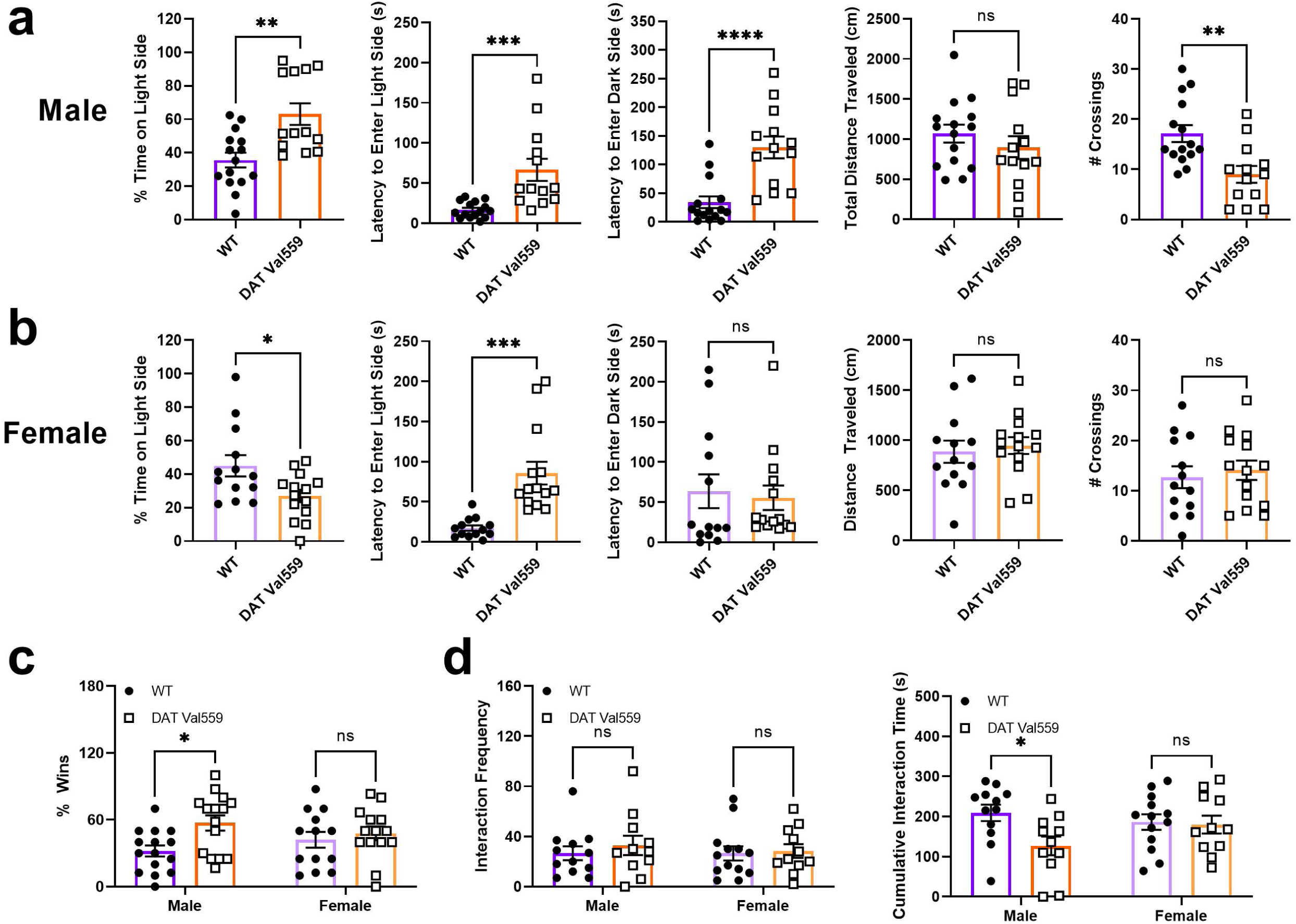
Threat aversion and social behavior in male and female DAT Val559 mice. The light/dark transition test was performed in WT male (n=15), WT female (n=13), DAT Val559 male (n=13) and DAT Val559 female (n=14) mice. The proportion of a 10 min recording session the animals spent on the light side, latencies to enter the light side and return to the dark side, distance traveled, and total number of light/dark transitions are depicted for **a** males and **b** females. **c** Tube test performance of WT male (n=15), WT female (n=13), DAT Val559 male (n=15) and DAT Val559 female (n=14) mice. Individual performance (% wins) is graphed. Male DAT Val559 mice won 79/123 total bouts (*P*=0.0001) and female DAT Val559 mice won 55/106 (*P*=0.68). **d** Social interaction was measured in WT male (n=12), DAT Val559 male (n=11), WT female (n=13) and DAT Val559 female (n=11) mice. Interaction time with the novel mouse as well as frequency of social approach are depicted. Data were analyzed by student’s t-test or two-way ANOVA with Sidak’s multiple comparisons test. ^*^*P*<0.05, ^**^*P*<0.01, ^***^*P*<0.001, ^****^*P*<0.0001. ns = not significantly different. Data are presented as mean ± SEM.

In the tube test, an assay that forces direct, but non-aggressive, social interaction, DAT Val559 males, but not females, displayed an increase in bout wins (Fig. 2c). However, in the three-chamber sociability assay, we noted that females of both genotypes and DAT Val559 males largely failed to exit the center chamber and explore either the novel object or the novel mouse (Fig. S7a), hindering efforts to assess social preference in DAT Val559 mutants. Thus, we moved on to a simplified social interaction test wherein a sex-matched novel mouse was enclosed in a cage within a single chamber. Here we found that male, but not female, DAT Val559 mice displayed a reduction in interaction time with the novel mouse though the total frequency of interaction was unaltered (Fig. 2d).

In the social interaction test, DAT Val559 females displayed a decrease in interaction with the empty cage during habituation (Fig. S7b), prompting direct interrogation of object recognition memory in our model. Although distance traveled and object exploration time did not differ significantly between groups (Fig. S7c), we noted a difference in novel object preference and discrimination in DAT Val559 females compared to WT littermates, a phenotype not observed in males (Fig. 3a). Further, while Male WT and DAT Val559 mice showed no relationship between engagement of novelty as a function of total distance traveled, female WT animals demonstrated a significant negative correlation in this measure. DAT Val559 females, conversely, assumed the pattern seen in males (Fig. 3b).

**Fig. 3.**
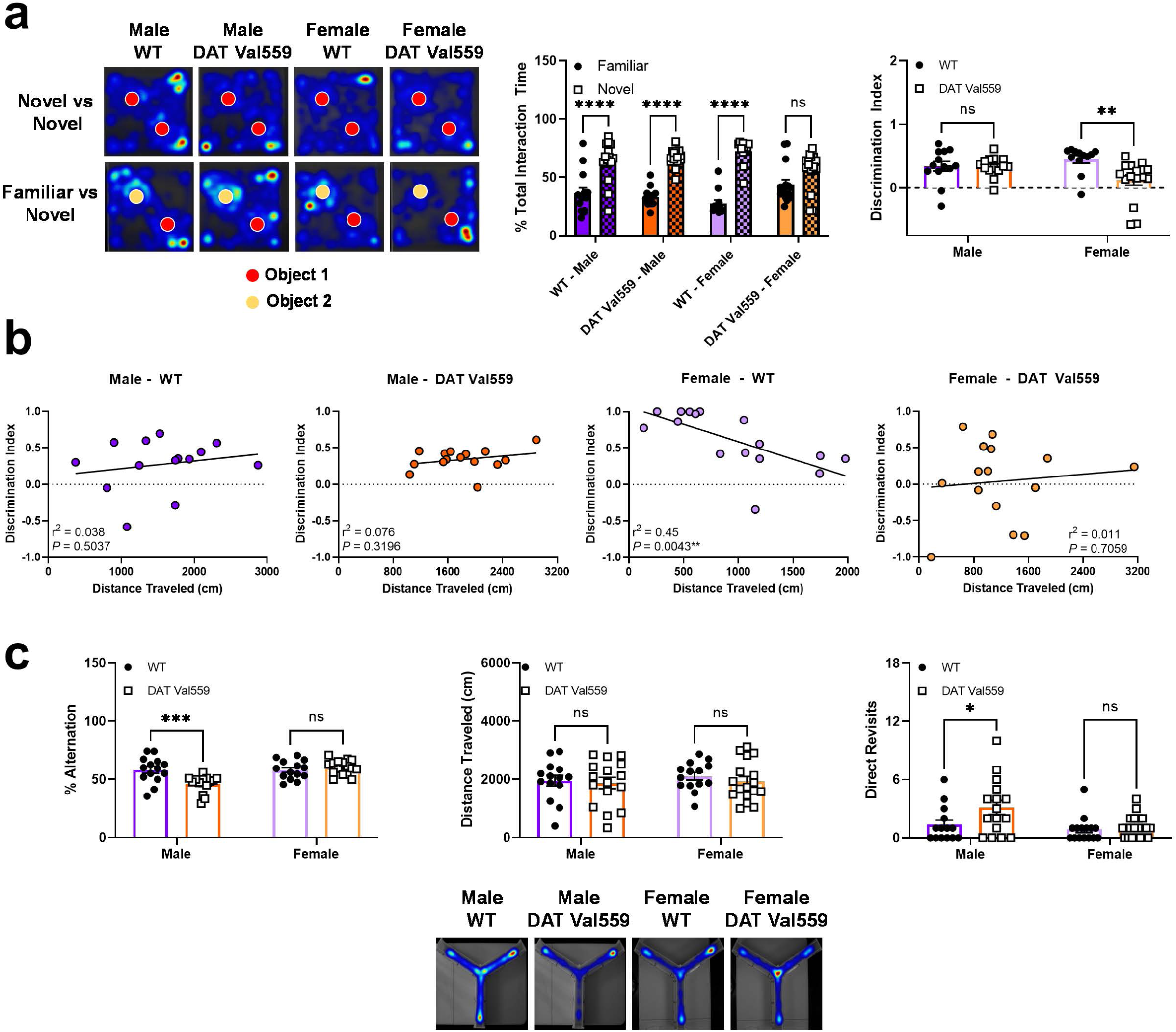
Male and female DAT Val559 mice display divergent deficits in memory tasks. **a** The novel object recognition test was performed with WT male (n=15), WT female (n=12), DAT Val559 male (n=15) and DAT Val559 female (n=16) mice. Total time spent interacting with a familiar vs novel object and object discrimination index are depicted along with representative heat maps for the training and testing days of the paradigm. **b** Correlation analysis in the novel object recognition task depicting the distance traveled and discrimination index for individual mice. A simple linear regression was fit to the data to determine goodness of fit (r^2^) and the degree to which the slope significantly deviates from 0 (*P*). **c** Y Maze spontaneous alternation test in WT male (n=14), WT female (n=14), DAT Val559 male (n=16) and DAT Val559 female (n=16) mice. % Alternations and distance traveled are shown along with representative heat maps for exploration patterns in the maze. Data were analyzed by two-way ANOVA with Sidak’s multiple comparisons test. ^*^ *P*<0.05, ^**^*P*<0.01, ^***^*P*<0.001, ^****^*P*<0.0001. ns = not significantly different. Data are presented as mean ± SEM.

Finally, to determine whether the female-specific DAT Val559 disruption seen in novel object discrimination extended to other forms of memory, we examined working memory in the Y maze spontaneous alternation task. We found a significant reduction in percent alternation that could not be attributed to hypoactivity in DAT Val559 males but not females (Fig. 3c). To consider whether male-specific deficits in alternations might arise from altered memory capacity versus an elaboration of repetitive behavior, we examined frequency of repetitive exploration of the same arm (direct revisits), and found males but not females demonstrated an increase in direct revisits (Fig. 3c).

### D2AR-DAT coupling reveals females-specific impact of DAT Val559 on transporter phosphorylation, trafficking and DA clearance in vStr

At this stage, our data were inconsistent with the hypothesis that female sex simply negates phenotypes exhibited by DAT Val559 male mice. Rather, the two sexes often displayed a unique profile of behavioral responses to the variant, raising the possibility that sex might dictate which, rather than whether, behavioral phenotypes are displayed. This idea led us to consider the possibility that sex might influence how different dopaminergic circuits regulate or respond to functional changes in DAT. In our prior work using male mice, we discovered that stimulation of D2ARs with the agonist quinpirole increases WT DAT Thr53 phosphorylation and enhances transporter surface trafficking in the dStr but not in the vStr^25^. Moreover, we found that, as a consequence of the anomalous DA efflux of DAT Val559^22^, D2ARs become constitutively activated in the dStr, thereby sustaining DAT Val559 phosphorylation and elevated surface expression of “leaky” DATs^25^. Surprisingly, in females, activation of D2Rs increased DAT Thr53 phosphorylation in the vStr but not the dStr (Fig. 4a,b). This same sex-dependent flip in the circuit dependence of D2AR regulation of DAT was evident with respect to D2AR-dependent DAT trafficking (Fig. 4c,d). Moreover, DAT Val559 phosphorylation and trafficking were elevated relative to WT in the absence of quinpirole and no further increase in these parameters was seen when quinpirole was added (Fig. 4a-d). Instead, quinpirole decreased the elevated surface trafficking of DAT Val559 in female vStr (Fig. 4d).

**Fig. 4.**
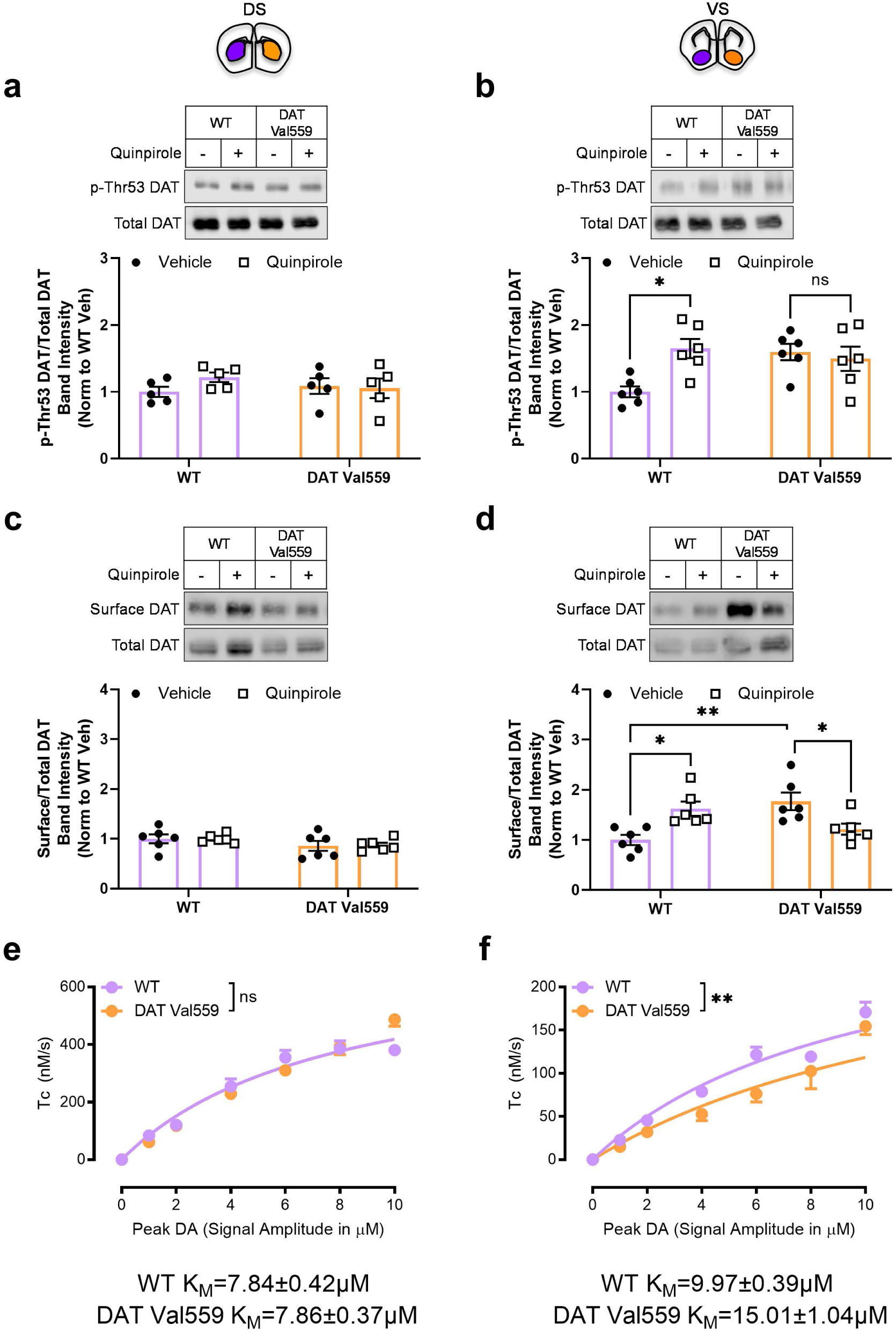
DAT Val559 induces elevated transporter surface trafficking and phosphorylation, but compromises uptake in the NAc of females. Basal and quinpirole (1 μM) driven DAT phosphorylation (pThr53) in slices containing **a** dStr and **b** NAc of WT (n=5) and DAT Val559 (n=5) females. Basal and quinpirole (1 μM) driven DAT surface expression in slices containing **c** dStr and **d** NAc of WT (n=6) and DAT Val559 (n=6) females. Surface and phospho-DAT levels were normalized to total DAT and data are expressed relative to WT vehicle. Dopamine clearance rates (T_C_) revelaed by *in vivo* high speed chronoamperometry in **e** dStr and **f** NAc of WT (n=20) and DAT Val559 (n=14-20) females. Curves conformed well to Michaelis–Menten kinetics in both WT (*r*^2^ dStr = 0.75, NAc = 0.86) and DAT Val559 (*r*^2^ dStr = 0.87, NAc = 0.69). K_M_ values are shown on the graphs. Data were analyzed by two-way ANOVA with Sidak’s multiple comparisons test. ^*^*P*<0.05, ^**^ *P*<0.01. ns = not significantly different. Data are presented as mean ± SEM.

Increased DAT Val559 surface expression triggered by tonic D2AR activation may overwhelm the capacity to clear evoked DA release due to preferential ADE. Indeed, we reported that in male DAT Val559 mice, clearance of pulse applied DA, as assessed by high-speed *in vivo* chronoamperometry, is significantly reduced in the dStr but not the vStr^25^. In keeping with our findings of elevated DAT Val559 Thr53 phosphorylation and surface trafficking, we observed a decreased *in vivo* DA clearance rate in the vStr, but not the dStr of DAT Val559 females (Fig. 3e,f). These findings demonstrate that the molecular and cellular impact of the DAT Val559 variant at native DA synapses *in vivo* parallels the switch in subregional action exhibited by D2AR-mediated transporter phosphorylation and trafficking.

### D2R inhibition rescues distinct behavioral phenotypes in male and female DAT Val559 mice

In males, application of the D2-type DA receptor antagonist sulpiride normalized elevated DAT Val559 surface trafficking in acute dStr brain slices, with a similar reversal of altered DA clearance evident *in vivo*^18, 25^. In male WT animals, sulpiride antagonized AMPH-induced locomotion (Fig. 5a, S8a, S8b). However, sulpiride failed to further attenuate AMPH-induced locomotion in DAT Val559 males. In females, where AMPH produces an equivalent rise in locomotor activation in both WT and DAT Val559 mice (Fig. 1b), sulpiride increased the locomotor response to AMPH in DAT Val559 mice, but produced the expected decrease in WTs (Fig. 5b, S8c, S8d)^32^. Although inhibition of D2Rs failed to rescue acute AMPH-dependent locomotor activation in male DAT Val559 mice, we were able to rescue behavioral phenotypes characteristic of DAT Val559 mice of each sex via blockade of D2Rs. In the NOR task, as noted above, female, but not male, DAT Val559 mice displayed a deficit in object discrimination. Whereas sulpiride failed to impact the performance of WT females in the NOR task, it improved object discrimination in DAT Val559 females (Fig. 5c) without influencing locomotor activity or object exploration time (Fig. S9). Conversely, DAT Val559 males, but not females, exhibited a reduction in spontaneous alteration in the Y maze accompanied by an increase in direct revisits that was also not seen in female mice. Again, sulpiride had no effect on the performance of WT males in the task, but restored spontaneous alternation and direct revisits in DAT Val559 males to levels observed in their WT littermates without influencing gross locomotion (Fig. 5d). Together, these findings indicate that tonic activation of D2Rs drives specific, sex-dependent behavioral phenotypes in DAT Val559 mice, which plausibly derive from sex-biased, region-specific D2AR-DAT coupling.

**Fig. 5.**
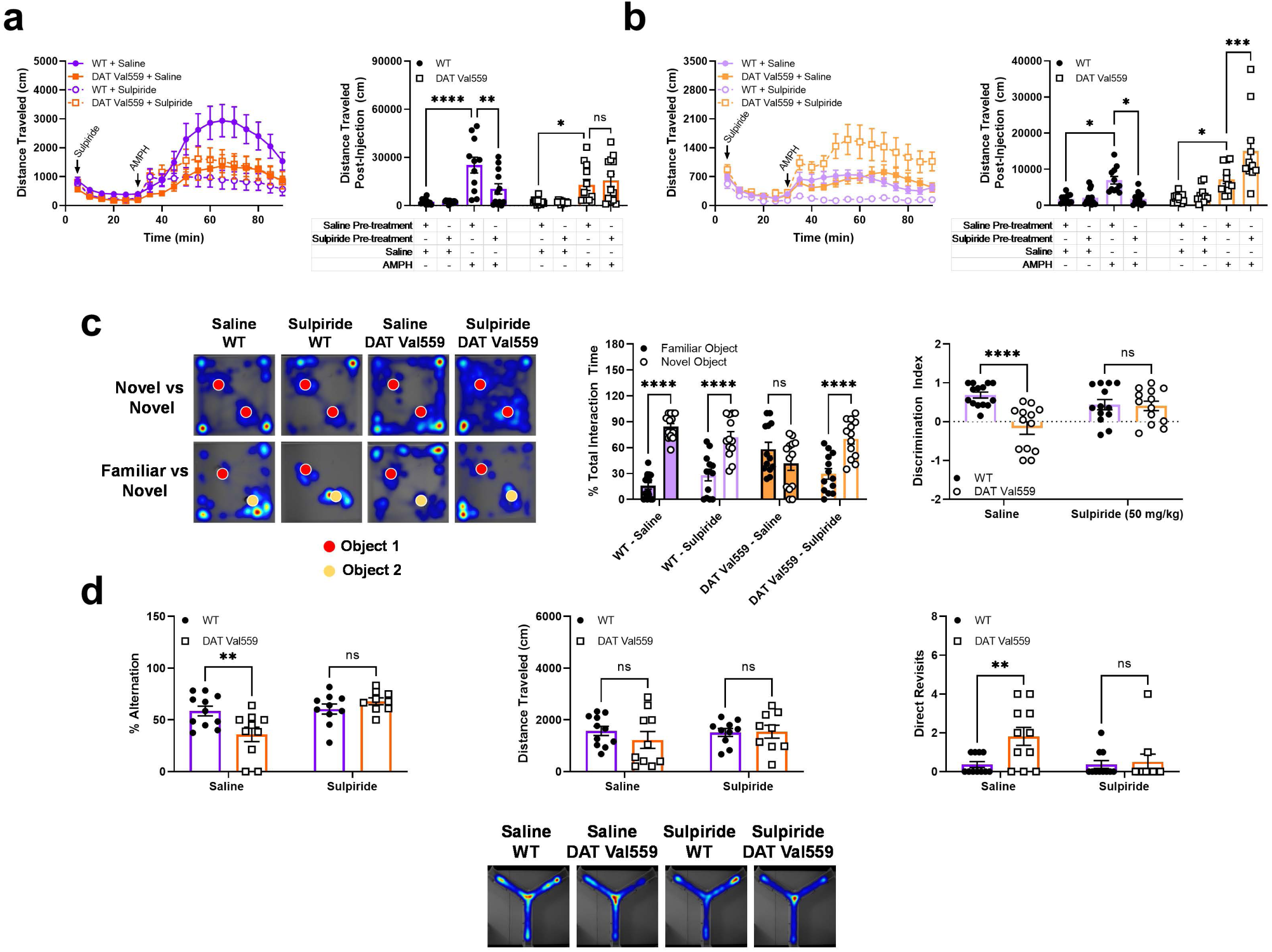
D2R antagonism rescues select behavioral phenotypes of DAT Val559 male and female mice. WT male (n=11-12), WT female (n=11-12), DAT Val559 male (n=13) and DAT Val559 female (n=11-12) were pre-exposed to the D2R antagonist sulpiride (50 mg/kg, i.p.) 30 mins prior to AMPH challenge (3 mg/kg, i.p.). Distance traveled (full time course data as well as summary data for the 60 min recording period post-drug injection) are depicted for **a** male and **b** female mice. **c** WT (n=13-14) and DAT Val559 (n=13-14) females were given a single injection of saline or sulpiride (50 mg/kg, i.p.) 30 mins prior to NOR testing on both the training and testing (days 3-4) days of the paradigm. Total time spent interacting with a familiar vs novel object and object discrimination index are depicted along with representative heat maps for the training and testing days of the paradigm. **d** WT (n=10-11) and DAT Val559 (n=9-10) mice were given saline or the D2 antagonist sulpiride (50 mg/kg, i.p.) 30 mins prior to Y maze testing. % Alternations and distance traveled are shown along with representative heat maps for exploration patterns in the maze. Data were analyzed by two-way ANOVA or two-way repeated measures ANOVA (time courses) with Sidak’s multiple comparisons test. ^*^*P*<0.05, ^**^*P*<0.01, ^***^*P*<0.001, ^****^*P*<0.0001. ns = not significantly different. Data are presented as mean ± SEM. **Fig. 5** DAT Val559 males exhibit D1R hyposensitivity and loss of AMPH-dependent ERK 1/2 activation. **a** Inhibition of specific DA uptake was assessed in whole striatal synaptosomes isolated from male and female WT (n=5) and DAT Val559 (n=6) mice exposed to increasing concentrations of AMPH (10^−9^ to 10^−4^ M). Nonlinear curves were fit to the data to determine IC_50_ values (WT male, 131 ± 26 nM; DAT Val559 male, 252 ± 55nM; WT female, 217 ± 30 nM; DAT Val559 female, 255 ± 30 nM). AMPH (3 mg/kg, i.p.)-induced **b** DA and **c** 5-HT elevations in the dStr measured using *in vivo* microdialysis. Eluates from WT and DAT Val559 male (n=6) and female (n=8-9) mice were pooled in 20 min bins and expressed as averages over time. **d** WT male (n=10), WT female (n=16), DAT Val559 male (n=9) and DAT Val559 female (n=15) were administered a single injection of the D1R agonist SKF83822 (0.5 mg/kg, i.p.). **e** Phosphorylation of ERK 1/2 was determined in whole striatal lysates from WT and DAT Val559 male (n=6) and female (n=6-7) mice following AMPH (3 mg/kg, i.p.) challenge (15 min). Representative immunoblots are shown. Phosphorylated ERK 1/2 was normalized to total protein and data are expressed relative to WT saline treated animals. The full time course data as well as summary data for the 60 min recording period post-drug injection are shown. Data were analyzed by two-way ANOVA or two-way repeated measures ANOVA (time courses) with Sidak’s multiple comparisons test. ^*^*P*<0.05, ^**^*P*<0.01, ^***^*P*<0.001,^****^*P*<0.0001. ns = not significantly different. Data are presented as mean ± SEM.

## Discussion

Here, aided by the DAT Val559 model, we unmask a sex-dependent alteration in a key, *in vivo* DAT regulatory mechanism: the ability of D2ARs to control DAT surface expression that governs high-affinity clearance of extracellular DA. This sex-dependence is not one of simple presence or absence, but rather, due to an underlying circuit specificity of D2AR control of DAT, is one of common impact on different pathways in males and females. We show that in males, coupling of D2ARs to DAT, as revealed in studies of D2-type agonist stimulation of DAT phosphorylation and trafficking, flexibly regulates DAT surface expression and DA clearance in the dStr projection fields of nigral DA neurons, a region heavily implicated in DA-driven locomotor behavior and habit formation^25^, but not the vStr terminal fields of VTA DA neurons. In contrast, in females, regulation of DAT by D2ARs is seen in the vStr but not the dStr. In the dStr of males and the vStr of females, the ADE of DAT Val559 drives tonic D2AR activation that results in elevated basal DAT phosphorylation and surface trafficking. Though we used a racemic sulpiride mixture containing both heteroreceptor-selective and non-selective enantiomers^33^, the restoration of spontaneous alteration and object recognition memory in DAT Val559 males and females, respectively, is consistent with a contribution of tonically active D2ARs to sex-biased behavioral phenotypes in DAT Val559 mice. Notably, quinpirole treatment resulted in a unique decrease in surface DAT levels in DAT Val559 females, a trend we previously observed in males^25^. This effect is consistent with prior reports indicating that D2 agonists such as aripiprazole can antagonize the actions of DA with high dopaminergic tone^34^, as we might expect with ADE-driven tonic activation of D2ARs in DAT Val559 mice. A similar effect was not observed for DAT phosphorylation suggesting that the latter modification may saturate at lower DA concentrations than D2AR-dependent DAT surface trafficking. Indeed, Foster et al. showed that Thr53 mutations impacted DAT-dependent efflux but not surface trafficking *in vitro*^13^.

Our behavioral studies reveal changes in both baseline and drug-stimulated behaviors that distinguish male and female DAT Val559 mice (summarized in Table S1). Based on our analysis of DA clearance *in vivo* (Fig. 3e,f)^25^, we expect that male and female DAT Val559 mice would exhibit deficits in behaviors associated with nigrostriatal vs mesolimbic DA activity, respectively. Knockdown of DAT in the SNc or optogenetic stimulation of SNc DA neurons produces social deficits^35^, observations consistent with male-specific alterations in sociability in the DAT Val559 model. Although direct stimulation of VTA neurons projecting to the NAc has been reported to promote social approach, it is unclear if the circuits driving social behavior are conserved in females^36^. DAT Val559 males display a similar frequency of social interaction, but a decrease in cumulative interaction time. These behaviors are not consistent with a lack of social approach, but rather a sort of social obliviousness wherein the DAT Val559 males more rapidly lose interest in the social stimulus.

The spontaneous alternation task engages circuitry involved in perception, attention, spatial memory, and also motivation and exploration, functions typically associated with DA action in the frontal cortex and, to a lesser degree, hippocampus^37^. Indeed, while lesioning of the NAc fails to impact spontaneous alternation^38, 39^, VTA lesions decrease alternation rates^40^ implicating VTA projections to extrastriatal regions in alternation behavior. However, DA clearance remains unaltered in the vStr of DAT Val559 males, a region heavily innervated by the VTA. Though evidence linking the dStr to spontaneous alternation is limited, lesions of the dStr do diminish alternation behaviors in rats^41^ suggesting that this phenotype in our DAT Val559 males, which we know is driven by D2R hyperactivity, could derive from D2R/DAT dysregulation in the nigrostriatal circuit.

The hypophagic effects of AMPH are blunted in DA-deficient mice, which lack tyrosine hydroxylase expression in all DAergic neurons^42, 43^, but can be recovered following restoration of DA signaling in the dStr^42^. DAT Val559 males, but not females, exhibited blunted hypophagia to AMPH. Thus, suppression of AMPH-induced hypophagia in DAT Val559 mice may arise from a sex-dependent penetrance of DAT Val559 and its regulation by D2ARs in the male but not female dStr. Similarly, while acquisition of CPP relies on DA release in the NAc, recent evidence suggests that the persistence of drug-context associations involves the medial PFC in males^44^. Our observation that the DAT Val559 mutation accelerated or delayed AMPH CPP in male or female mice, respectively, emphasizes that sex differences in DA signaling may also occur in the mesocortical circuit, which we have yet to investigate due to the sparse nature of DA projections to this region.

Our observation that DAT Val559 males, but not females, exhibit alterations in psychostimulant-dependent locomotor activation is also consistent with a selective perturbation by DAT Val559 in nigrostriatal projections. However, acute blockade of D2AR feedback with the antagonist sulpiride failed to restore AMPH responses in DAT Val559 males. Possibly, blunted AMPH-dependent locomotor activation in DAT Val559 males may result from desensitization of post-synaptic D1Rs due to tonic ADE as direct stimulation of D1Rs in DAT Val559 males fails to elicit a full locomotor response. While mice lacking DAT display a loss of D1R expression^6,45^, D1R binding is intact in striatal membranes from male DAT Val559 mice^23^. Rather, loss of D1R-dependent locomotor stimulation in DAT Val559 males may result from uncoupling of select effectors such as the kinase cascade leading to phosphorylation of ERK 1/2, activated by AMPH in D1R-expressing striatal medium spiny neurons^46, 47^ and known to mediate AMPH-dependent hyperactivity^48, 49^.

The pattern of behavior for DAT Val559 males in the light/dark transition test, characterized by increased latencies to travel between the light/dark boxes and a decrease in total crossings, is perhaps more consistent with pure disinhibition of exploratory drive rather than a lack of active threat avoidance. Female DAT Val559s, in contrast, display a classic anxiety phenotype with a decrease in time spent in the illuminated box and an increased latency to exit the dark that aligns well with literature indicating that catecholaminergic denervation of the amygdala decreases light side occupancy^50^. Importantly, dopaminergic projections that terminate in the amygdala originate in the VTA^51^ along with those to the ventral striatum, where we observe tonic D2AR-dependent feedback onto pre-synaptic DAT in DAT Val559 females.

Inhibition of D1 receptors in the perirhinal cortex, a critical mediator of memory consolidation^52, 53^, impairs performance in the NOR task^54^. VTA dopaminergic projections also innervate the rhinal cortex^51^. Thus, the deficit in NOR discrimination displayed by DAT Val559 females is consistent with disrupted function of VTA dopaminergic neurons, a conclusion that compels the implementation of projection specific manipulations of D2ARs. Nonetheless, our observation that NOR performance in DAT Val559 females can be improved with sulpiride provides supportive evidence that tonic D2AR activation by DAT Val559 in mutant females disrupts NOR. Though more frequently demonstrated in rats and humans, evidence indicates that females display superior performance in the object recognition task particularly when the novel and familiar objects are similar^55^. Indeed, our dataset highlights that, while WT males and females both successfully identify the novel object, the cognitive strategy employed^56^ may differ by sex with females appearing to achieve high discriminative capacity by maintaining focus on objects. Although inhibition of D2Rs clearly rescued NOR memory in DAT Val559 females, the correlation between activity and discrimination index was not restored (Fig. S10), indicating that D2R-independent mechanisms contribute to sex-dependent NOR strategies.

The neural substrate(s) underlying circuit-specific and sex-biased D2AR signaling could result from either ingrained, sex chromosome-derived variation or the impact of circulating sex hormones. Mediators of sex-dependence of D2AR-dependent DAT regulation could involve molecules that stabilize or limit receptor-transporter physical interactions^57^, that anchor these proteins into a critical microdomain^58^, or that participate in the signaling pathway that supports functional coupling of receptor and transporter such as ERK1 and PKCβ^29, 59^. Notably, DAT Thr53 phosphorylation has been reported to be elevated in estrus females^10^ alluding to possible direct modulation of DAT activity by estradiol and/or progesterone, though recent studies have described sex-dependent differences in DA release/reuptake dynamics across striatal subregions in slice preparations irrespective of estrus stage^14^.

Alterations in DA signaling have long been associated with multiple neuropsychiatric disorders, including ADHD, ASD, schizophrenia and substance abuse/addiction. In ADHD, noncoding genetic variation in *SLC6A3*, the gene encoding DAT, and DRD2 and other DA receptors have been associated with risk for ADHD though not in all studies^60-62^. Rather than considering risk per se, often studied with genomic markers that lack direct functional relevance, we^19, 63^ and others^64-69^ have sought to identify rare coding variation impacting DAT functional properties, efforts that have revealed intrinsic, circuit and behavior-level changes when modeled *in vitro* in transfected cells and *in vivo* when studied in transgenic mice and flies. We have pursued the DAT Val559 variant due to its support of ADE, a process akin to the actions of AMPH, which is supported by constitutive actions of D2Rs *in vitro*^70^ and *in vivo*^25^. Our current study extends our prior work in males, suggesting that efforts to characterize females with other mutants will likely be profitable. The DAT Val559 model and other DAT mutant mice also provide opportunities to identify molecular changes (e.g. transcriptome, epigenome, proteome) that drive sex-differences in DA-linked traits, including response to therapeutics.

## Conclusions

A “resilience” framework is often used to explain discrepancies in the sex bias observed in neuropsychiatric disorders. However, recent evidence suggests that sex bias can be due, at least in part, to differences in symptomology, morbidity, and associated comorbidities^71^ and the resultant failure of our diagnostic instruments to assure identification of the same disorder in both sexes. Our studies with DAT Val559 mice reinforce the critical importance of sex as a determining factor in the neurochemical, neurophysiological and behavioral consequences of altered DAT-dependent DA signaling. We show that not only do different DA projections exhibit major differences in DAT regulation by D2ARs, but also that these circuit differences differ by sex such that the behaviors arising from their genetic or pharmacological perturbations are distinct in males and females. The broader implications of our findings are two-fold. First, our work provides a clear example of how changes in behavioral traits thought to define distinct neurobehavioral disorders may have arisen from shared underlying causes, consistent with findings of shared polygenic risk factors for categorically distinct psychiatric disease^72^. Second, sexual dimorphism in the architecture of the DA system likely also influences the efficacy of treatment strategies aimed at reversing or mitigating disruptions in DA homeostasis in men and women. Revision of diagnostic criteria for psychiatric disease away from behavioral features and toward biological definitions, with recognition of the modulatory influence of sex, will be essential to tailor therapeutic interventions.

## Acknowledgements

The authors wish to acknowledge the infrastructure support from Blakely laboratory staff members Matthew Gross, Alaina Tillman and Zayna Gichi. The authors thank the Vanderbilt and Florida Atlantic University Mouse Neurobehavioral Cores for support of behavioral experiments and neurochemical analyses. Financial support for this work derives from the Postdoctoral Training Program in Functional Neurogenomics MH065215 (AS), NARSAD Young Investigator Grant from the BBRF (AS), a Max Kade fellowship (FPM), and NIH awards MH107132 (GLD) and MH086530 (RDB). RP and AB received grant support from the Office for Undergraduate Research and Inquiry (OURI) at FAU.

