## Supplementary Material for "Sex and Circuit Specific Dopamine Transporter Regulation Underlies Unique Behavioral Trajectories of Functional *SLC6A3* Coding Variation"

**Data supplement**

#### Authors/Affiliations

Adele Stewart PhD[1,3], Felix P. Mayer PhD[1], Raajaram Gowrishankar PhD[1], Gwynne L. Davis PhD[1], Lorena B. Areal PhD[1], Paul J. Gresch PhD[1,3], Rania M. Katamish[1], Rodeania Peart[2], Samantha E. Stilley[1], Keeley Spiess[1], Maximilian J. Rabil[1], Faakhira A. Diljohn[2], Angelica E. Wiggins[1], Roxanne A. Vaughan PhD[4], Maureen K. Hahn PhD[1,3], Randy D. Blakely PhD[1,3,*]

[1]Department of Biomedical Science, [2]Wilkes Honors College, and [3]Brain Institute, Florida Atlantic University, Jupiter, FL. [4]Department of Biomedical Sciences, University of North Dakota School of Medicine and Health Sciences, Grand Forks, North Dakota.

#### Supplemental Methods

*Striatal microdialysis*

Mice were anesthetized with isoflurane (5% for induction, 2% for maintenance during surgery) and placed in a stereotaxic frame. A guide cannula (CMA7) was placed 1 mm above the dStr (−0.86 AP from Bregma, ±1.6 mL and −2.0 DV from dura) and secured to the skull using glass ionomer cement (Instech Solomon, Plymouth Meeting, PA). After recovery from surgery (18–24 h), animals were placed in individual dialysis chambers (clear cylindrical enclosure, 14 cm diameter, 22 cm high; Instech Solomon). A microdialysis probe (CMA Microdialysis, Holliston, MA) with an active length of 2 mm was inserted into the guide cannula. One end of a tether was attached to the headpiece and the other end was attached to a liquid swivel (Instech Solomon) that was mounted on a counterbalanced arm above the dialysis chamber. The probe was then perfused with aCSF at a flow rate of 1.0 μL/min overnight. Approximately, 18-24 hours after surgery samples were collected every 20 min including 4 baseline samples and an additional 6 following drug injection. Dialysate samples were stored at −80 °C and analyzed by HPLC-EC for DA and 5-HT levels as described previously[25]. After each dialysis session, animals were overdosed with sodium pentobarbital, brains were removed and postfixed in 4% paraformaldehyde in PBS, sectioned, and then inspected for acceptable probe placement.

*Synaptosomal [$^3$H]DA uptake inhibition*

Brains were harvested following rapid decapitation and whole striata dissected on a pre-chilled metal platform. Synaptosomes were prepared from the striata of WT or DAT Val559 animals as previously described[31]. 25 μg of striatal synaptosome preparations were pre-incubated with amphetamine [$10^{-9}$-$10^{-4}$ M] for 10 min at 37°C prior to the addition of 20 nM [$^3$H]DA (specific activity 46 Ci/mmol; Perkin-Elmer, Waltham, MA). Control samples were incubated with assay buffer only, and non-specific uptake was determined in parallel samples to which 1 μM GBR-12909 (Sigma) was added. Assays were terminated by rapid filtration over 0.3%

polyethyleneimine-soaked GF/B glass microfiber filters (Whatman, Maidstone, UK) and washed three times with ice-cold PBS. Filters were placed into scintillation vials with 7 mL of Ecoscint H (National Diagnostics, Atlanta, GA) scintillation fluid, shaken overnight at RT, and radioactivity was quantified by using a TriCarb 2900TR scintillation counter (Perkin-Elmer). Assays were performed in triplicate for all conditions, averaged and expressed relative to control.

*Light/Dark Transition Test*

Anxiety-like behavior was determined in WT and DAT Val559 mice by exploiting the conflict in mice between spontaneous exploration and their aversion to brightly illuminated spaces[32]. An insert divided Med Associates locomotor chambers into 2 equally sized spaces, one brightly illuminated and one dark, linked by a door. Animals were placed in the dark chamber and allowed to freely explore both chambers for 10 min. The latency to enter the dark, latency to return to the light side, total time spent on each side of the apparatus, number of transitions and total distance traveled were recorded for each animal. So as not to disturb the animal's circadian rhythms, light/dark testing was performed during the inactive phase.

*Social Interaction Test*

Social interaction (SI) was assessed in an open field arena (40 cm x 40 cm x 40 cm) containing a clear mouse holder that allows for nose contact through the bars while preventing fighting, as previously described[33, 34]. First, test mice were allowed to habituate to the arena and the empty mouse holder for 10 min. Next, a visitor mouse was introduced to the test mouse by being placed in the holder and the animals were allowed to interact for 5 min. Mice used as visitors were stranger, same sex and age-matched heterozygous animals. SI was measured using Noldus Ethovision XT software (v13.0) to calculate the amount of time and frequency that mice spent exploring the visitor mouse in a predefined interaction zone.

*Tube Test*

Social interactions between WT and DAT Val559 mice were evaluated in the tube test as previously described[35]. The apparatus is a 30-cm-long, 2.7-cm-diameter clear acrylic tube with small acrylic funnels added to each end to facilitate entry into the tube. On two separate days before testing, each mouse was exposed to the tube, with progress through the tube resulting in the mouse being returned to the home cage. Mice that were resistant to entering the tube were gently encouraged to run with a gentle tail pull. Animals that failed to progress (taking >1 min, freezing or backing out) were removed from the test cohort. For tube test bouts, one WT and one DAT Val559 age, sex and approximate weight matched mice from different home cages were placed at the opposite ends of the tube and released. A subject was declared a "winner" when its opponent backed out of the tube. Each mouse was tested against 3-5 individuals from other cages and trials were repeated with each mouse beginning at either end to avoid position bias resulting in 6-10 bouts per individual. If neither animal backed out of the tube after a time period of 2 min, a draw was declared. Draws were excluded from analysis.

*Y Maze Test of Spontaneous Alternation*

Testing occurred in a Y-shaped maze combining 3 white, opaque plastic arms (14 × 4.5 × 40 cm) each at a 120°C angle from the others. Each subject was introduced to the center of the maze and allowed to freely explore all arms for 10 min. Locomotor activity and arm entries were monitored using Noldus Ethovision XT video tracking software (v13.0) and the percent alteration calculated as the number of alternations (incidences where animals visited each arm in turn without repeating an arm) divided by the total number of arm visit triads X 100. The number of incidences where an animal displayed consecutive visits to the same arm (direct revisits) was also monitored as a gross metric of perseverative behavior. A separate cohort of animals was given a single injection of saline or sulpiride (50 mg/kg, i.p.) 30 mins prior to Y maze testing.

Heatmaps tracking center point movement over the testing period were generated with Noldus Ethovision XT (v13.0) software.

*Novel Object Recognition (NOR) Test*

Object recognition memory was assessed utilizing a previously published protocol[36]. Briefly, on days 1 and 2 animals were habituated to the testing apparatus (40 cm x 40 cm x 40 cm square arena with opaque walls) for 10 min. On the training day (day 3) two identical objects (50 ml conical tubes containing Drierite™) were introduced to the chamber on the diagonal (at opposite corners) and animals were allowed to explore both objects for 10 min. On the testing day a novel object (T-25 flask filled with Drierite™ and tagged with a plastic triangle on the cap) replaced one familiar object and mice were again allowed to explore for 10 min. Object exploration was monitored with Noldus Ethovision XT software (v13.0). An animal was considered to be interacting with the object if the nose point detection fell within 2.5 cm of the object. The placement of the novel object was counterbalanced between subjects to preclude the introduction of spatial bias. An object discrimination index was calculated for each subject by subtracting the time spent exploring the familiar object from the time spent exploring the novel object and dividing by the total time exploring both objects. A separate cohort of animals was given a single injection of saline or sulpiride (50 mg/kg, i.p.) 30 mins prior to testing on object exploration days 3 and 4. Heatmaps tracking nose point movement over the testing period were generated with Noldus Ethovision XT (v13.0) software.

*Three Chamber Sociability Test*

Social behavior was evaluated in a three-chamber polycarbonate apparatus with 4-inch sliding gates separating the 7 × 9-inch chambers. Each subject was first allowed to acclimate to the apparatus with free access to all chambers during a 10-min habituation. A stimulus mouse was then placed in a small, rectangular plastic cage in one side chamber and a clean empty cage introduced in the opposite side chamber. The stimulus mouse was an adult WT animal of the same sex that had been previously habituated to the cage in three one-hour sessions across 3 days. Stimulus mice that exhibited excessive climbing or biting of the cage bars were excluded. The subjects were allowed to explore all chambers for 10 mins. Time spent in each chamber and the total distance traveled was determined using Noldus (Leesburg, VA) Ethovision XT video tracking software (v13.0).

### Supplemental Figures

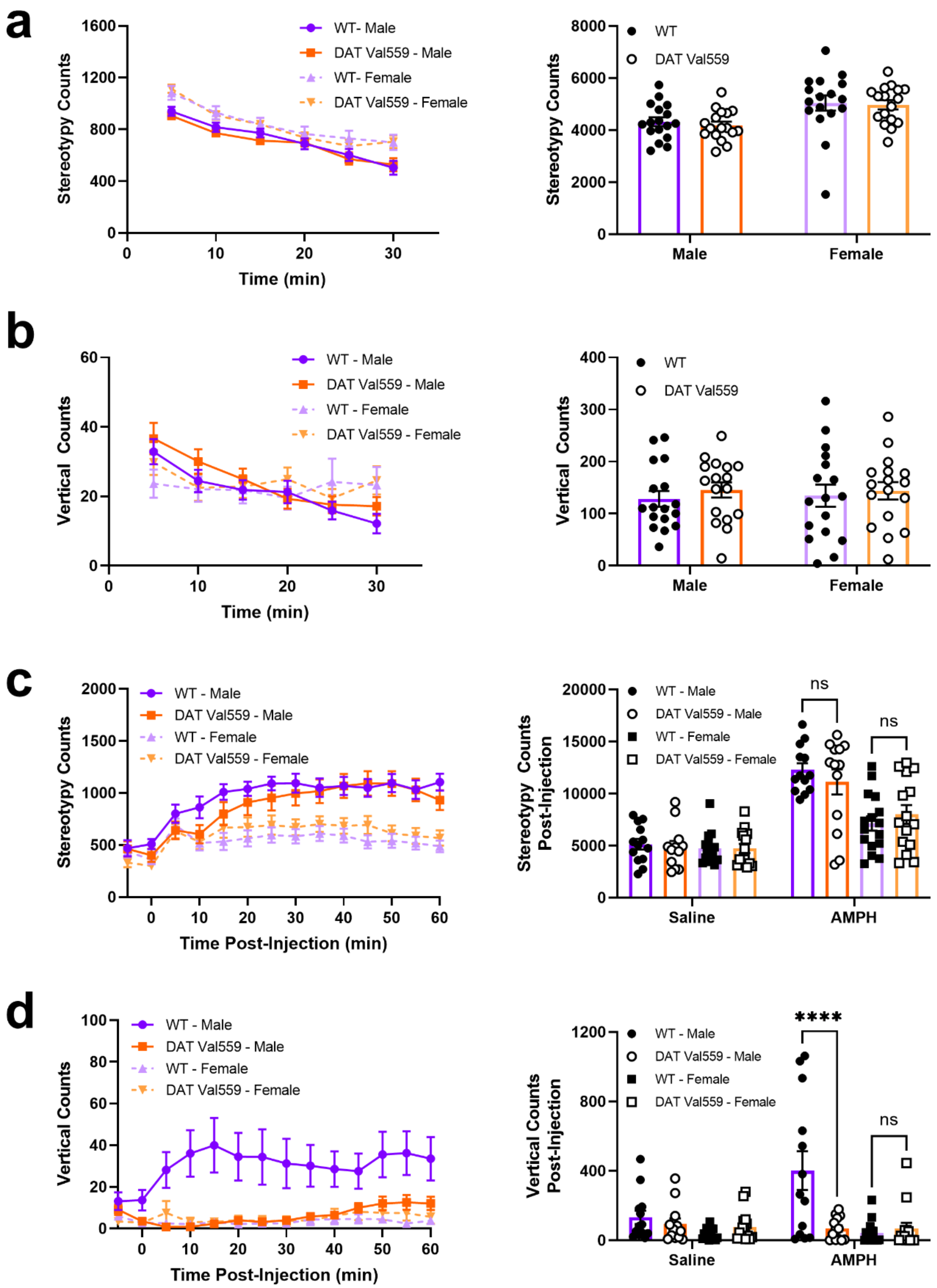

**Fig. S1** DAT Val559 males exhibit blunted AMPH-dependent vertical locomotor activation. WT (n=17) and DAT Val559 (n=17) male and females were allowed to freely explore an open field chamber for 30 mins. **a** Stereotypy and **b** rearing were monitored over time and summarized for the complete recording period. Acute AMPH (3 mg/kg, i.p.)-driven stereotypy and vertical locomotion in WT male (n=13), WT female (n=16), DAT Val559 male (n=13) and DAT Val559 female (n=15) mice. **c** Stereotypy and **d** rearing were monitored over time and summarized for the 60 minutes post-drug injection. Data are presented as mean ± SEM.

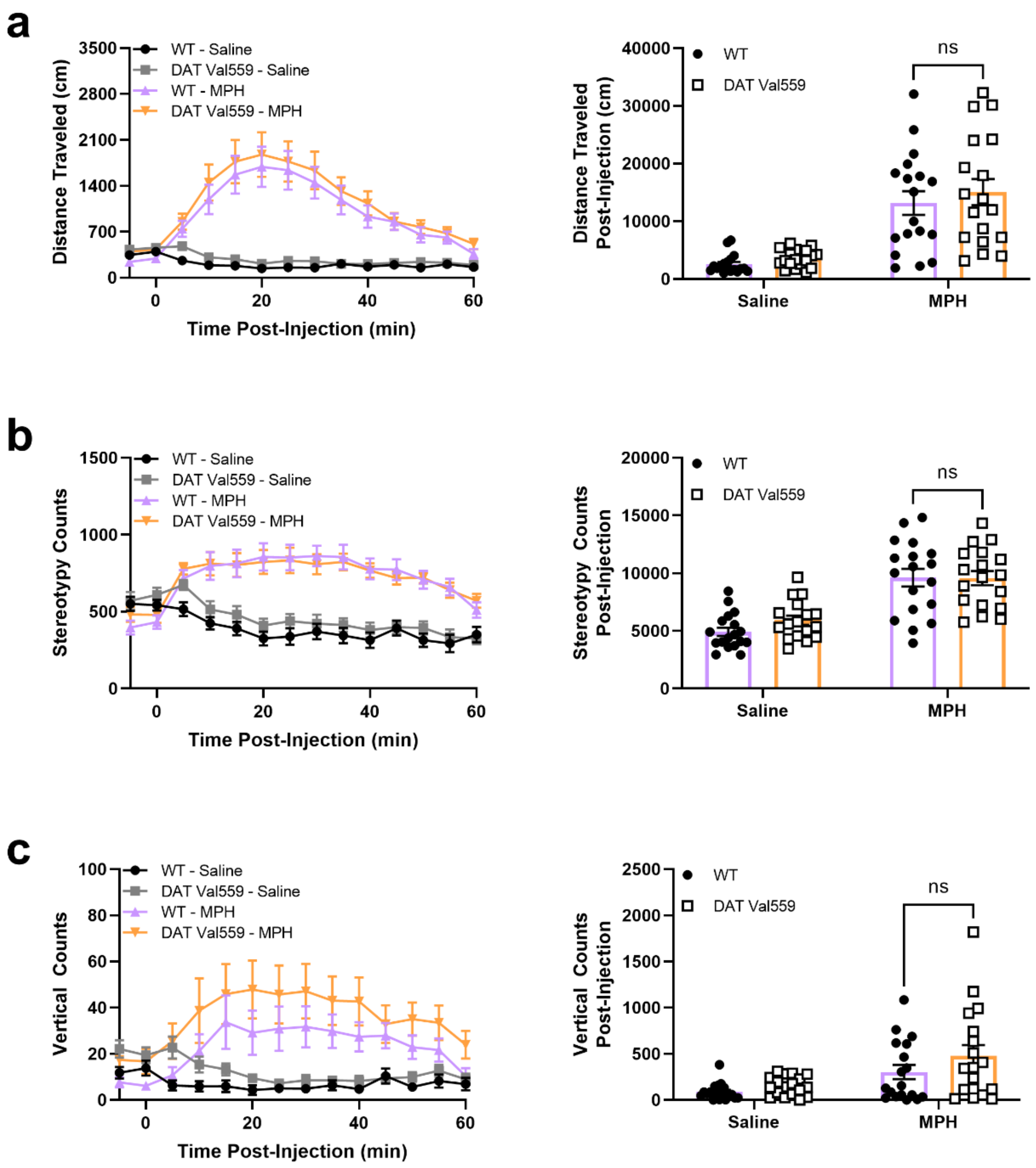


**Fig. S2** Female DAT Val559 display normal locomotor response to MPH. WT (n=18) and DAT Val559 (n=18) females were given a single MPH (10 mg/kg, i.p.) injection and **a** horizontal distance traveled, **b** stereotypy, and **c** vertical locomotion monitored overtime. Figures **a-c** present both full time course data as well as summary data for the 60 min recording period post-drug injection. Data were analyzed by two-way ANOVA with Sidak's multiple comparisons test. ns = not significantly different. Data are presented as mean ± SEM.

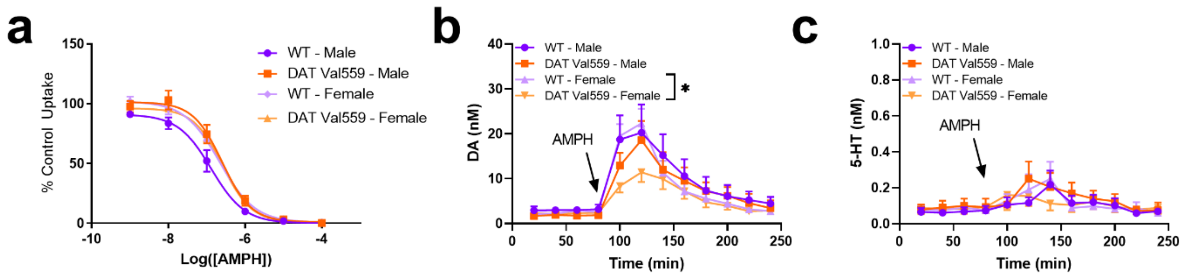


**Fig. S3** AMPH inhibits DA uptake and elevates DA in the dStr of male DAT Val559 mice. **a** Inhibition of specific DA uptake was assessed in whole striatal synaptosomes isolated from male and female WT (n=5) and DAT Val559 (n=6) mice exposed to increasing concentrations of AMPH ($10^{-9}$ to $10^{-4}$ M). Nonlinear curves were fit to the data to determine $IC_{50}$ values (WT male, 131 ± 26 nM; DAT Val559 male, 252 ± 55nM; WT female, 217 ± 30 nM; DAT Val559 female, 255 ± 30 nM). AMPH (3 mg/kg, i.p.)-induced **b** DA and **c** 5-HT elevations in the dStr measured using *in vivo* microdialysis. Eluates from WT and DAT Val559 male (n=6) and female (n=8-9) mice were pooled in 20 min bins and expressed as averages over time. Data were analyzed by two-way repeated measures ANOVA (time courses) with Sidak's multiple comparisons test. *$P$<0.05. Data are presented as mean ± SEM.

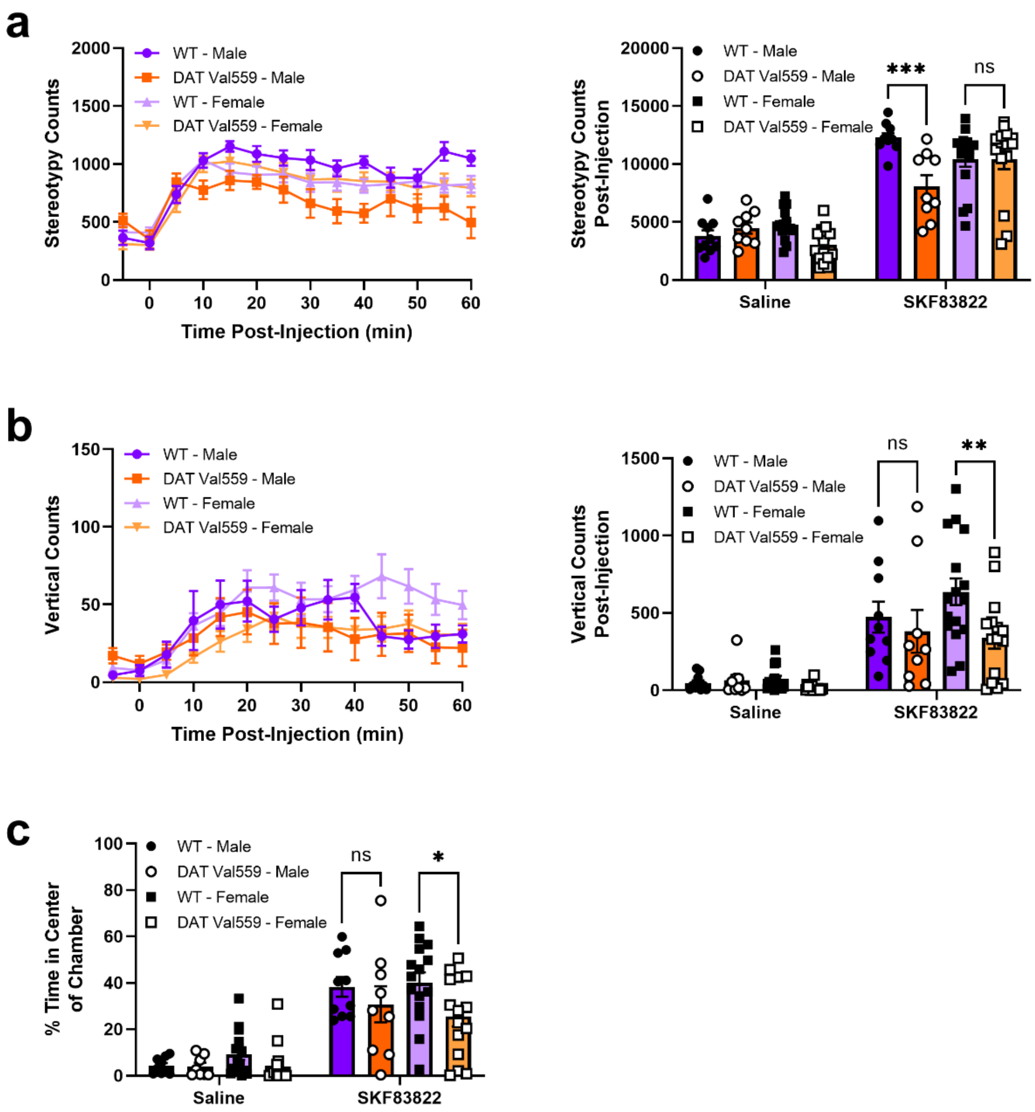


**Fig. S4** D1R-dependent locomotor behaviors differ by genotype and sex in WT and DAT Val559 mice. **a** Stereotypy, **b** rearing, and **c** center occupancy in WT male (n=10), WT female (n=16), DAT Val559 male (n=9) and DAT Val559 female (n=15) mice following a single injection of the D1R agonist SKF83822 (0.5 mg/kg, i.p.). Time course data are presented along summary data for the 60 min recording period post-drug injection. Data were analyzed by two-way ANOVA with Sidak's multiple comparisons test. **P*<0.05, ***P*<0.01, ****P*<0.001. ns = not significantly different. Data are presented as mean ± SEM

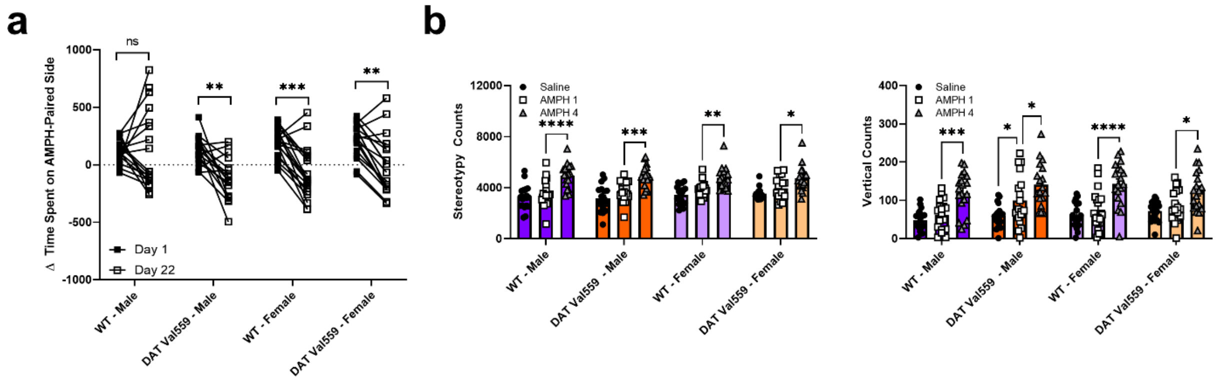


**Fig. S5** Supplemental AMPH CPP data. AMPH (3 mg/kg, i.p.) CPP was performed in WT male (n=18), WT female (n=19), DAT Val559 male (n=18) and DAT Val559 female (n=19) mice. **a** Trajectory of CPP score on from day 1 to day 22 post-conditioning. **b** Locomotor sensitization for stereotypy and rearing. Data were analyzed by two-way ANOVA with Sidak's multiple comparisons test. **P*<0.05, ***P*<0.01, ****P*<0.001, *****P*<0.0001. ns = not significantly different. Data are presented as mean ± SEM.

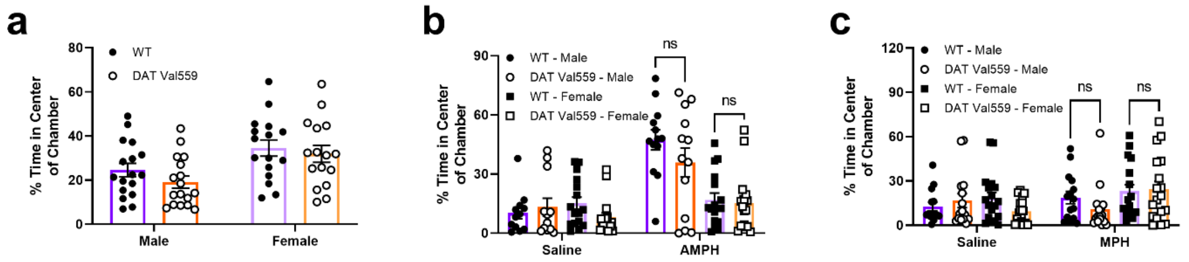


**Fig. S6** Baseline and psychostimulant-driven thigmotaxis in the open field. **a** WT (n=17) and DAT Val559 (n=17) male and females were allowed to freely explore an open field chamber for 30 mins and the proportion of total time spent in the center of the chamber is depicted. **b** Acute AMPH (3 mg/kg, i.p.)-driven center occupancy in WT male (n=13), WT female (n=16), DAT Val559 male (n=13) and DAT Val559 female (n=15) mice. Summary data for the 60 min recording period post-drug injection is provided. **c** WT (n=18) and DAT Val559 (n=18) females and WT (n=15) and DAT Val559 (n=16) males were given a single MPH (10 mg/kg, i.p.) injection. Center occupancy for the 60 min recording period post-drug injection is shown. Data were analyzed by two-way ANOVA with Sidak's multiple comparisons test. ns = not significantly different. Data are presented as mean ± SEM.

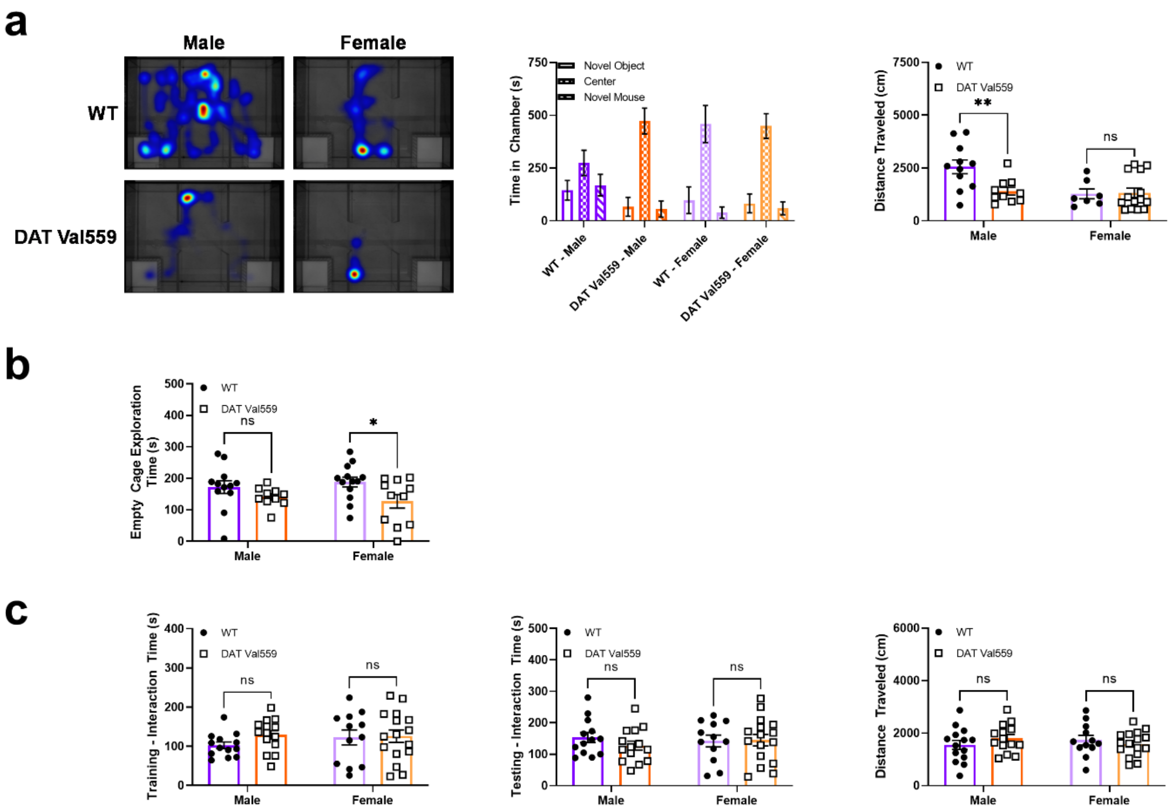
a
Male
Female
WT
DAT Val559
Novel Object
Center
Novel Mouse
Time in Chamber (s)
750
500
250
0
WT - Male
DAT Val559 - Male
WT - Female
DAT Val559 - Female
Distance Traveled (cm)
7500
5000
2500
0
WT
DAT Val559
**
ns
Male
Female
b
Empty Cage Exploration Time (s)
500
400
300
200
100
0
WT
DAT Val559
ns
*
Male
Female
c
Training - Interaction Time (s)
400
300
200
100
0
WT
DAT Val559
ns
ns
Male
Female
Testing - Interaction Time (s)
500
400
300
200
100
0
WT
DAT Val559
ns
ns
Male
Female
Distance Traveled (cm)
6000
4000
2000
0
WT
DAT Val559
ns
ns
Male
Female

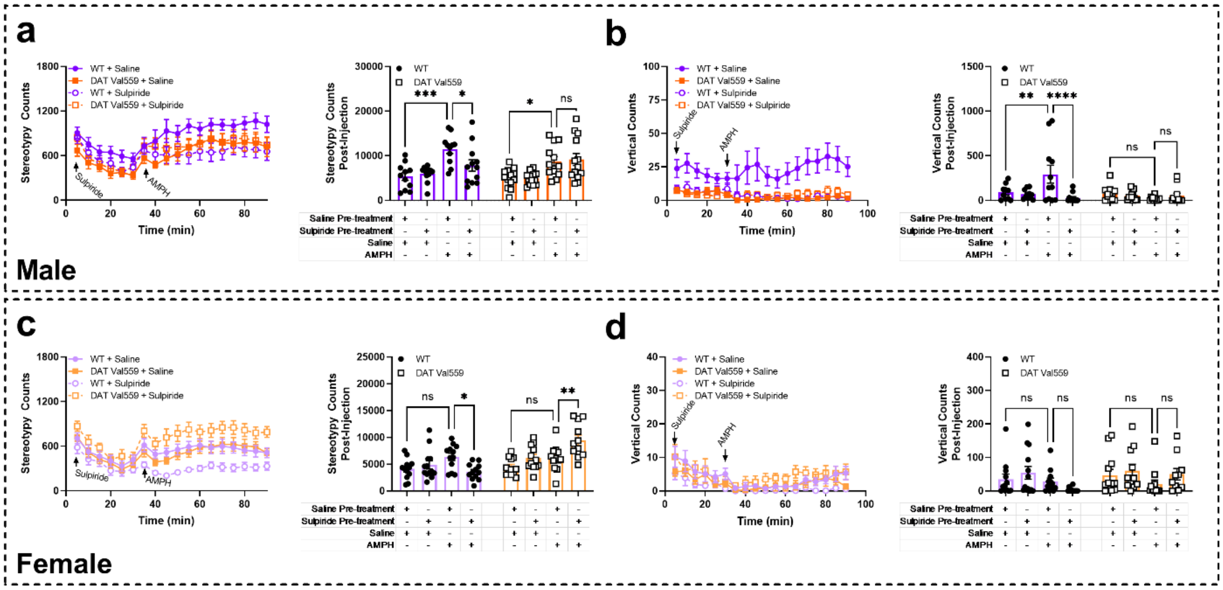


**Fig. S8** Sex and DAT Val559 genotype differentially influence the impact of D2R antagonism on AMPH-induced stereotypy and rearing. WT male (n=11-12), WT female (n=11-12), DAT Val559 male (n=13) and DAT Val559 female (n=11-12) were pre-exposed to the D2R antagonist sulpiride (50 mg/kg, i.p.) 30 mins prior to AMPH challenge (3 mg/kg, i.p.). **a** Stereotypy and **b** rearing in males. **c** Stereotypy and **d** rearing in females. Data are presented as both full time course data as well as summary data for the 60 min recording period post-drug injection. Data were analyzed by two-way ANOVA or two-way repeated measures ANOVA (time courses) with Sidak's multiple comparisons test. **P*<0.05, ***P*<0.01, ****P*<0.001, *****P*<0.0001. ns = not significantly different. Data are presented as mean ± SEM.

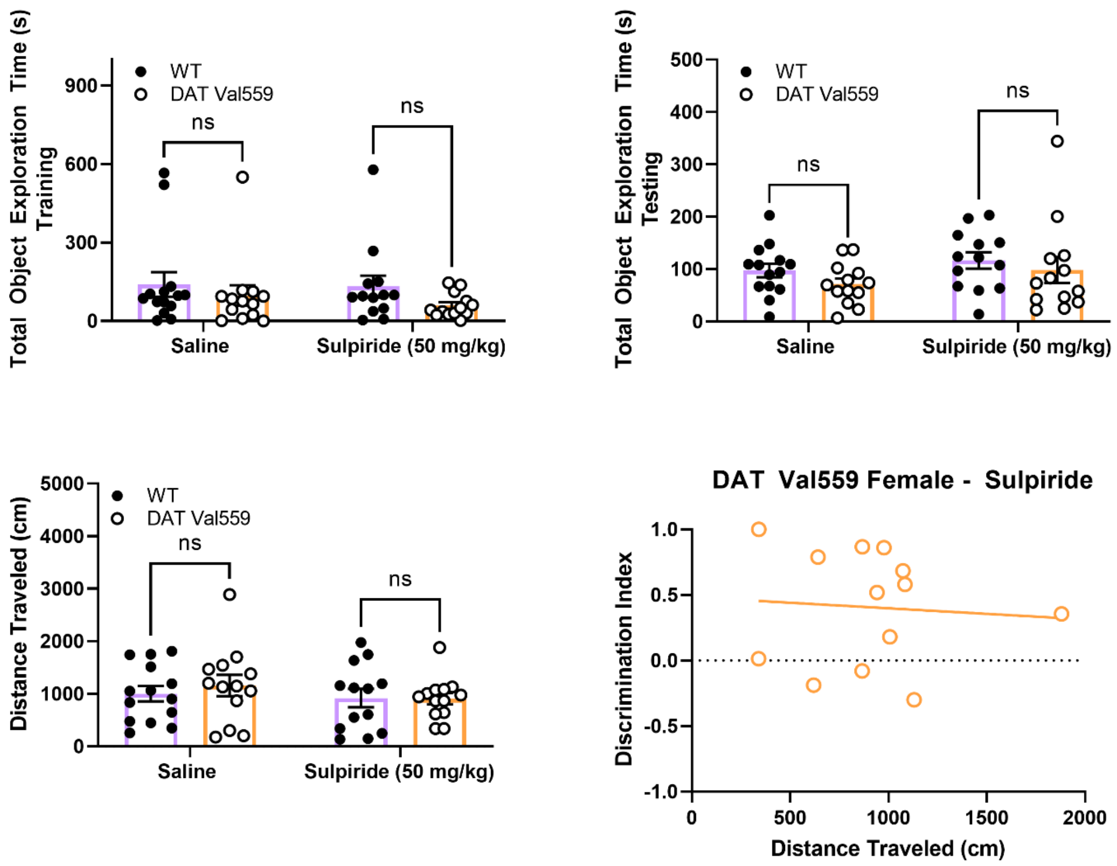


**Fig. S9** Supplemental data for the novel object recognition sulpiride rescue experiment. WT (n=13-14) and DAT Val559 (n=13-14) females were given a single injection of saline or sulpiride (50 mg/kg, i.p.) 30 mins prior to NOR testing on both the training and testing (days 3-4) of the paradigm. Total distance traveled during the testing phase as well as total object exploration time during both the training and testing phases are shown. The correlation between discrimination index and distance traveled is also provided for sulpiride-treated DAT Val559 females. ns = not significantly different. Data are presented as mean ± SEM.

**Table S1**: Summary of behavioral phenotypes in male and female DAT Val559 mice. Experimental paradigms where are differences was detected are highlight in bold. Male only phenotypes are coded blue; female only phenotypes are pink; and phenotypes shared by both sexes are purple.

| | Phenotype of DAT Val559 Mice | |
|---|---|---|
| | **Male** | **Female** |
| *Acute Psychostimulant Action* | | |
| Amphetamine (AMPH) Locomotion | **Blunted** | Normal |
| Methylphenidate (MPH) Locomotion | **Blunted** | Normal |
| SKF83822 Locomotion | **Blunted (Horizontal/Stereotypy)** | **Blunted (Rearing/Center Occupancy)** |
| *Chronic Psychostimulant Action* | | |
| AMPH CPP - Acquisition | Normal | Normal |
| AMPH - CPP Extinction | **Accelerated** | **Delayed** |
| AMPH - Locomotor Sensitization | Normal | Normal |
| AMPH - Weight Loss | **Blunted** | Normal |
| *Baseline Behaviors* | | |
| Locomotion | Normal | Normal |
| Light/Dark Transition Test | **↑ Light Occupancy** | **↓ Light Occupancy** |
| Tube Test | **↑ Wins** | Normal |
| Social Interaction Test | **↓ Interaction** | Normal |
| Novel Object Recognition | Normal | **↓ Discrimination** |
| Y Maze Spontaneous Alternation | **↓ Alternation** | Normal |
